# A GPCR screening in human keratinocytes identifies that the metabolite receptor HCAR3 controls epithelial proliferation, migration, and cellular respiration

**DOI:** 10.1101/2023.05.30.542853

**Authors:** M. Pilar Pedro, Katherine Lund, Sun Woo Sophie Kang, Ting Chen, Christina H. Stuelten, Natalie Porat-Shliom, Ramiro Iglesias-Bartolome

**Author notes:** **Correspondence**: Ramiro Iglesias-Bartolome, PhD LCMB/CCR, National Cancer Institute, 37 Convent Drive, Building 37 Room 2050C, Bethesda, MD 20892.

## Abstract

Epithelial cells in the skin and other tissues rely on signals from their environment to maintain homeostasis and respond to injury, and G protein-coupled receptors (GPCRs) play a critical role in this communication. A better understanding of the GPCRs expressed in epithelial cells will contribute to understanding the relationship between cells and their niche and could lead to developing new therapies to modulate cell fate. This study used human primary keratinocytes as a model to investigate the specific GPCRs regulating epithelial cell proliferation and differentiation. We identified three key receptors, hydroxycarboxylic acid-receptor 3 (HCAR3), leukotriene B4-receptor 1 (LTB4R), and G Protein-Coupled Receptor 137 (GPR137) and found that knockdown of these receptors led to changes in numerous gene networks that are important for maintaining cell identity and promoting proliferation while inhibiting differentiation. Our study also revealed that the metabolite receptor HCAR3 regulates keratinocyte migration and cellular metabolism. Knockdown of HCAR3 led to reduced keratinocyte migration and respiration, which could be attributed to altered metabolite use and aberrant mitochondrial morphology caused by the absence of the receptor. This study contributes to understanding the complex interplay between GPCR signaling and epithelial cell fate decisions.

## Introduction

The skin relies on self-renewing keratinocyte stem cells to replenish lost cells and heal wounds, making it a valuable system for studying cell fate decisions (1–3). The fate of basal keratinocytes includes either proliferation, to generate stem cells and transient-amplifying cells, or differentiation, which maintains the multilayer epidermis and its barrier function. The balance between proliferation and differentiation is crucial for proper tissue homeostasis, and its dysregulation can lead to aging-related disorders, impaired tissue regeneration, and cancer (4–6).

Keratinocytes must adequately sense their microenvironment to coordinate cell fate decisions, respond to stress, and restore homeostasis. G-protein-coupled receptors (GPCRs), the largest family of membrane signal transducers (7), are capable of sensing multiple extracellular molecules, including hormones, inflammatory mediators, nutrients, and metabolites, and can regulate cell fate in stem cells (8, 9). Understanding the GPCR-regulated signaling pathways in skin biology will shed light on the mechanisms stem cells use to sense and adapt to environmental changes. Moreover, given that more than one-third of all therapeutic drugs target GPCRs (10), identifying receptors involved in epithelial cell fate regulation can facilitate the development of pharmacological interventions to increase tissue regenerative capacity and modulate pathological conditions.

GPCRs relay their signaling through heterotrimeric G protein-dependent and independent pathways and are categorized into five classes: A (rhodopsin), B (secretin), C (glutamate), Frizzled, and Adhesion. Analysis of gene expression in mouse epidermis has shown that multiple GPCRs are expressed in keratinocytes, and the function of some of these receptors in the regulation of skin biology is well known (11). However, the functions of many GPCRs remain unclear due to their genetic and signaling redundancy and broad expression in multiple cell types. In addition, studies on GPCR pathways usually employ mouse models, highlighting the need for further investigation into the specific effects of GPCRs in human cells.

This study aimed to advance our understanding of epidermal cell fate regulation by GPCRs by characterizing the endogenous receptors expressed in human primary keratinocytes. Our findings reveal that hydroxycarboxylic acid-receptor 3 (HCAR3), leukotriene B4-receptor 1 (LTB4R), G Protein-Coupled Receptor 153 (GPR153), and G Protein-Coupled Receptor 137 (GPR137), play essential roles in modulating human keratinocyte proliferation and differentiation through the regulation of diverse gene expression networks. Overexpression of HCAR3, LTB4R, and GPR137, but not GPR153, resulted in aberrant development and differentiation of keratinocytes in 3D organotypic cultures. We also discovered that HCAR3, an understudied receptor exclusive to higher primates, is involved in keratinocyte migration and metabolism. Remarkably, HCAR3 regulates oxidative phosphorylation and fatty acid oxidation and is required to sustain proper mitochondrial energy output in keratinocytes. Our study provides a valuable resource for characterizing GPCRs in epithelial cells and sheds light on the potential role of cellular metabolism in keratinocyte proliferation, differentiation, and migration.

## Results

### Profiling GPCRs in human keratinocytes

To gain a better understanding of the GPCR landscape in keratinocytes, we conducted RNA sequencing (RNAseq) analysis on primary human epidermal keratinocytes (HEK) under non-differentiating and calcium-induced differentiation conditions (Fig. 1A). Our analysis revealed high expression levels of several GPCRs known to regulate skin biology (Fig. 1B). Of note, the β2 adrenergic receptor (*ADRB2*) has a role in wound closure in mice (12, 13); *LTB4R* is involved in allergic skin inflammation (14); sphingosine-1-phosphate receptors (*S1PR*) ligand is involved in skin hyperplasia (15); the F2R Like Trypsin Receptor 1 (*F2RL1*, also known as PAR-2) is involved in skin hyperplasia and squamous cell carcinogenesis (16); and the lysophosphatidic acid receptors (*LPAR*) ligand induces keratinocyte proliferation, skin hyperplasia and wound healing (17, 18).

**Figure 1:**
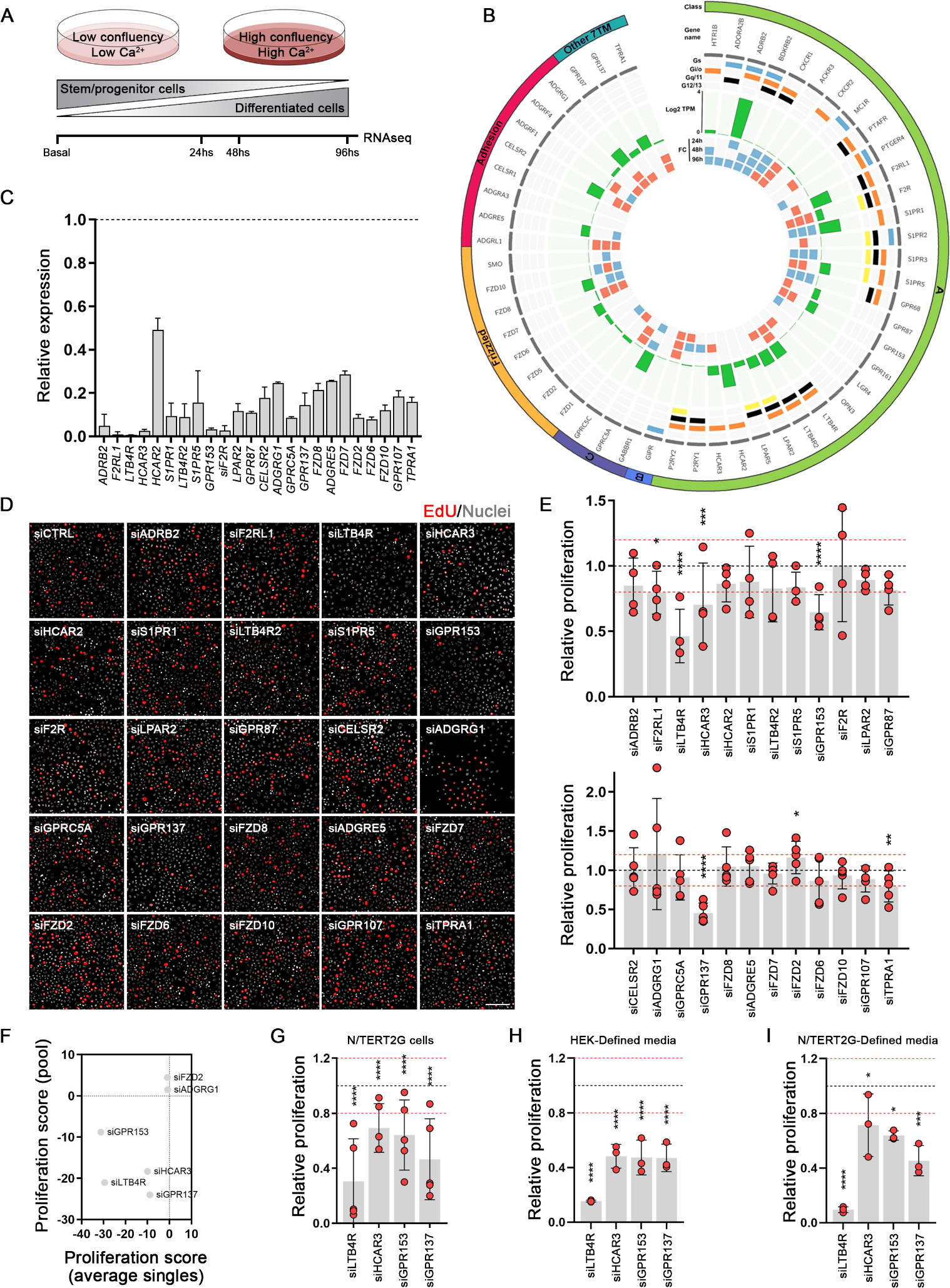
GPCRs regulating human keratinocyte proliferation. **(A)** Schematic representation of RNA-seq analysis of human skin keratinocytes (HEK) under basal and differentiated conditions. **(B)** Circos plot representing the GPCR expression in HEK cells showing class, gene name, Gα-coupling, level of expression (log2 of transcripts per million, Log2TPM), and fold change (FC) over basal/non-differentiated cells. Blue and red squares represent down-(q<0.05, FC≤-1.5) or up-(q<0.05, FC≥1.5) regulation, respectively. GPCRs with expression levels of TPM≥0.5 are shown. **(C)** qRT-PCR analysis showing the expression of the selected GPCRs in HEK cells 72 h after transfection with the corresponding siRNA for that gene. Values are indicated as fold over siCTRL (non-targeting control), shown as a black dotted line. **(D)** Representative images of EdU staining of siCTRL and siGPCRs transfected cells. Nuclei staining is shown in white, and EdU-positive cells are colored red. Scale bar: 100 μm. **(E)** Quantification of EdU-proliferation assay in primary HEK cells, 72hs after siRNA transfection with the indicated siRNAs (Class A top panel, non-Class A bottom panel). Results are normalized to non-targeting siRNA, represented as a black dotted line. Values of ±20% change in proliferation are shown as red dotted lines. n=4-5 independent experiments. **(F)** Proliferation scores comparing individual versus pool siRNA experiments (calculated as indicated in the methods section). **(G)** Quantification of EdU-proliferation assays in N/TERT2G cells presented as in E (n=5 independent experiments). **(H-I)** Quantification of EdU-proliferation assays in HEK (H) or N/TERT2G (I) cells grown in defined media and presented as in E (n=3 independent experiments). In E, G, H, and I: no asterisk p>0.05; *p<0.05; **p<0.01; ***p< 0.001; ****p< 0.0001; 2-way ANOVA followed by t-test.

We screened a set of the 24 highest expressed receptors with a pooled-small interference RNA (siRNA) library to identify the GPCRs affecting human keratinocyte proliferation (Fig. 1C). Knockdown of LTB4R, HCAR3, GPR153, and GPR137 significantly decreased primary HEK proliferation by more than 20% compared to non-targeting siRNA (Fig. 1D and E). LTB4R and GPR137 also reduced the proliferation of differentiated keratinocytes (Fig. S1A). Notably, the adhesion receptor ADGRG1 (also known as GPR56) and Frizzled Class Receptor 2 (FZD2) significantly increased the proliferation of differentiated HEK (Fig. S1A).

To confirm and validate the pooled siRNA results, we used four individual siRNAs to knockdown the identified GPCRs (Fig. S1B). Knockdown of LTB4R, HCAR3, GPR153, and GPR137 significantly reduced keratinocyte proliferation, while the efficacy of ADGRG1 and FZD2 siRNAs was inconsistent, suggesting possible off-target effects (Fig. 1F and S1C). Moreover, LTB4R, HCAR3, GPR153, and GPR137 knockdown reduced proliferation of immortalized human N/TERT2G keratinocytes (Fig. 1G, S1D and E). siRNAs for the GPCRs reduced proliferation in both full media (containing bovine pituitary extract, Fig. 1E and G) and defined media (Fig. 1H, 1I), indicating that the effects were not dependent on potential GPCR ligands present in the pituitary extract. Analysis of single cell data from human skin (19) confirmed the expression of *LTB4R*, *HCAR3*, *GPR153*, and *GPR137* in basal and differentiated (suprabasal) keratinocytes (Fig. S1F). Based on our results, we focused on LTB4R (also known as BLT1), HCAR3 (also known as GPR109B or HCA3), and the orphan receptors GPR153 and GPR137. Both LTB4R and HCAR3 are Gαi coupled receptors, highlighting the significance of this pathway in regulating keratinocyte cell growth (20).

### HCAR3, LTB4R, and GPR137 modulate transcriptional networks involved in keratinocyte proliferation and differentiation

To elucidate the mechanisms and signaling pathways responsible for the observed phenotypes, we performed RNAseq in primary HEK transfected with pooled siRNAs for LTB4R, HCAR3, GPR153, GPR137, and non-targeting sequence control. Principal component analysis (PCA) and Pearson correlation of the global RNA levels showed differential clustering of the siGPCRs with respect to control samples (Fig. S2A and B). Examination of differentially regulated transcripts revealed that GPCRs affected the expression of numerous genes (Fig. 2A). Gene ontology (GO) analysis of biological processes showed that 234 overlapping genes were involved in cell cycle processes (Fig. 2B), emphasizing the roles of the receptors in regulating keratinocyte proliferation.

**Figure 2:**
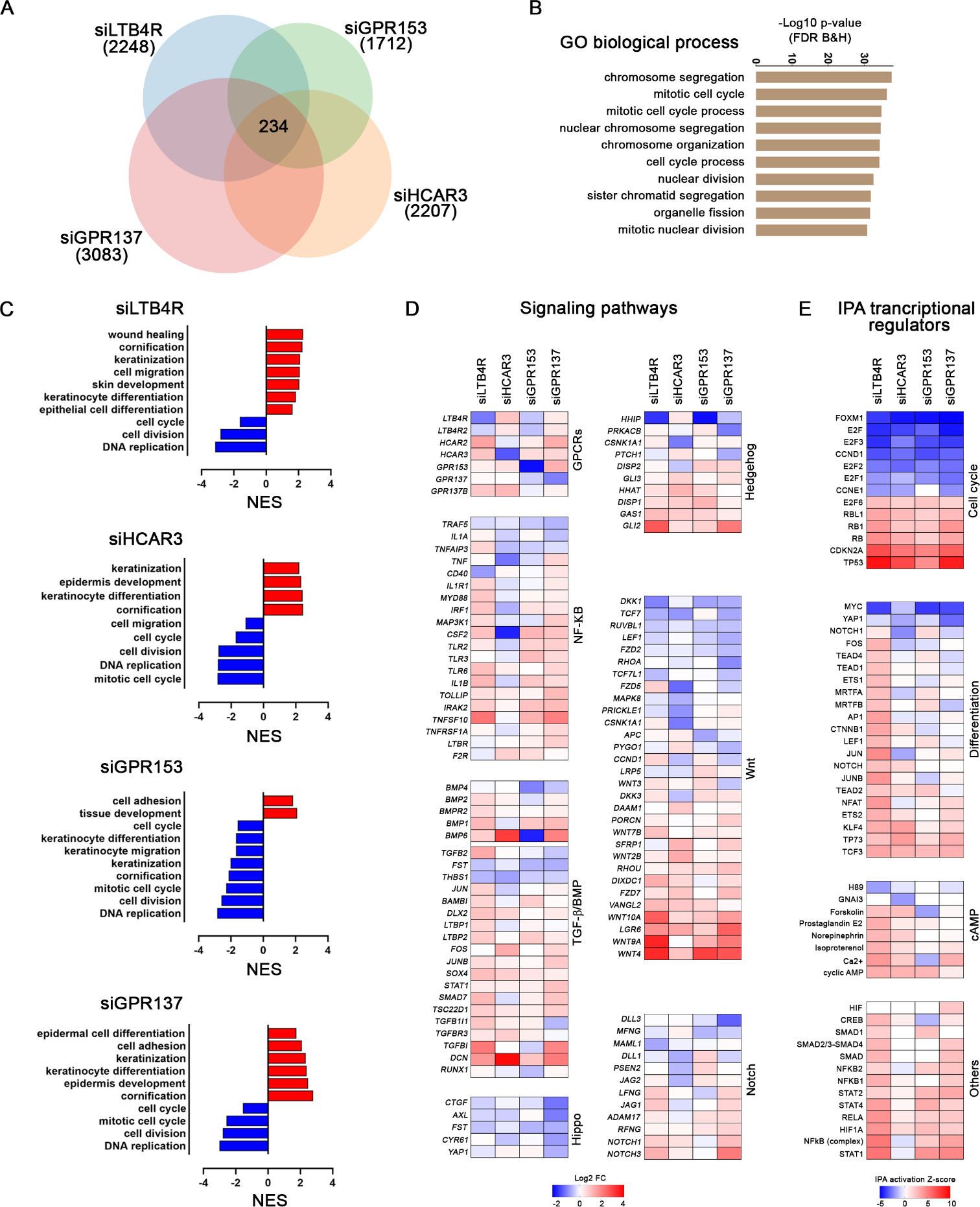
GPCRs control proliferation and differentiation gene networks. **(A)** Venn diagram showing the number of genes up- and down-regulated (q<0.05, FC≥±1.5) in RNAseq analysis of siGPCRs transfected cells. Common differentially regulated genes for all the siGPCRs are indicated in the middle (234). **(B)** Graph indicating the gene ontology (GO) biological process terms enriched in common differentially regulated genes. **(C)** Gene set enrichment analysis showing the normalized enrichment score (NES) for selected GO biological processes in each GPCR dataset. **(D)** Graph indicating expression fold change (Log2 FC) of genes related to canonical signaling pathways in each of the siGPCRs datasets. **(E)** IPA functional analysis of the activation state of upstream regulators with the indicated cell functions.

GO with normalized enrichment scores (NES) of biological processes in each dataset indicated that knockdown of LTB4R, HCAR3, and GPR137 activated differentiation and skin development, while downregulated processes related to cell cycle and DNA replication (Fig. 1C and). The knockdown of GPR153 downregulated both keratinocyte differentiation and cell cycle networks (Fig. 1C). Consistent with these findings, the siRNAs for LTB4R, HCAR3, and GPR137, but not GPR153 upregulated genes associated with all stages of keratinocyte differentiation (Fig. S2C). Western blot analysis for the differentiation markers KLF4 and keratin 10 (K10) confirmed the activation of differentiation programs following the knockdown of LTB4R, HCAR3, and GPR137 (Fig. S2D).

All four GPCRs showed common gene regulation trends on key epidermal pathways (Fig. 2D). Knockdown of LTB4R, HCAR3, GPR153, and GPR137 resulted in reduced Hippo signaling targets and differential regulation of NFKB, Wnt, Notch, and Hedgehog genes (Fig. 2D). Ingenuity pathway analysis (IPA) of transcriptional regulator networks confirmed that cell cycle-related networks, MYC and YAP1, are particularly affected by the knockdown of these GPCRs (Fig. 2E). Cyclic AMP (cAMP)-related gene networks were differentially regulated by LTB4R and HCAR3 (Fig. 2E), possibly due to reduced Gαi signaling resulting from the knockdown of these receptors.

Our gene expression analysis confirms the regulation of cell cycle gene networks by LTB4R, HCAR3, GPR153, and GPR137 and reveals numerous pathways affected by their downregulation. To further investigate these GPCRs, we generated doxycycline (Dox) inducible (i) N/TERT2G cells that overexpress LTB4R, HCAR3, GPR153, and GPR137 (iLTB4R, iHCAR3, iGPR153, and iGPR137; Fig. S3A). We found that overexpression of all four GPCRs increased AKT phosphorylation, while HCAR3 and GPR137 also activated ERK phosphorylation (Fig. 3A). In addition, HCAR3 induced activation of the mTOR target pS6 and reduced pCREB (Fig. 3A), a target for cAMP and protein kinase A (PKA).

**Figure 3:**
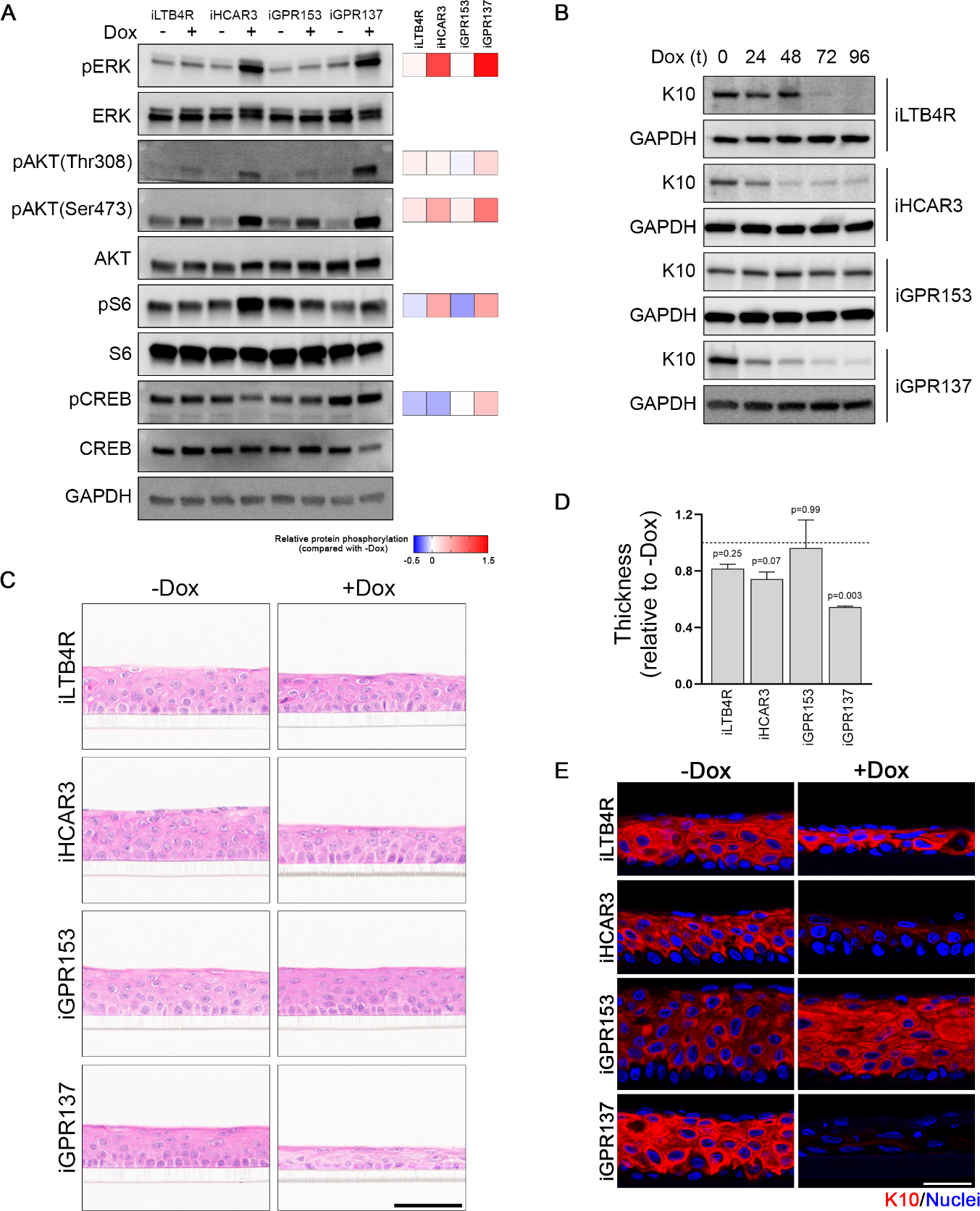
GPCR overexpression affects proliferation/differentiation balance and skin development. **(A)** Western blot showing expression of the indicated markers in cells treated (+) or not (-) with doxycycline (Dox) for 96 h to induce expression in inducible (i) GPCR stable cell lines. Relative levels for the phosphorylated bands normalized to the respective total protein and divided by the-Dox control is shown to the right. **(B)** Western blot showing the expression of keratin 10 (K10) in cells treated as indicated in A. **(C)** Hematoxylin & Eosin staining of organotypic assays without (-Dox) or with (+Dox) doxycycline to induce expression of the indicated GPCRs. Scale bar: 60µm. **(D)** Quantification of organotypic cultures thickness shown in C. n=3 independent experiments performed in duplicates; one-way ANOVA followed by t-test. **(E)** Representative images of immunofluorescence experiments showing the expression of keratin 10 (K10) in red and nuclei in blue in the organotypic cultures presented in C. Scale bar: 30 µm.

Confirming the effect of the GPCRs in keratinocyte differentiation, we observed a reduction of K10 in a time-dependent manner upon expression of LTB4R, HCAR3, and GPR137, but not GPR153 (Fig. 3B). HCAR3 and GPR137 reduced K10 expression even under differentiating conditions (Fig. S3B). In 3D skin-organotypic cultures, we observed abnormal development upon expressing LTB4R, HCAR3, and GPR137, while GPR153 did not have an effect (Fig. 3C and D). LTB4R, HCAR3, and GPR137 reduced differentiated cells as labeled by K10 expression (Fig. 3E), indicating that the reduction in skin thickness upon overexpression of the GPCRs could be due to aberrant differentiation. These findings suggest that GPCR-regulated gene networks play a critical role in human keratinocyte proliferation and differentiation and that their downregulation or overexpression can lead to altered epidermal homeostasis.

### HCAR3 regulates keratinocyte migration

Our RNAseq analysis revealed that GPCR knockdown led to differential regulation of gene networks related to keratinocyte migration, particularly in siHCAR3 cells (Fig. 2C). Given the crucial role of migration in skin wound healing, we conducted scratch-wound assays to examine the effect of LTB4R, HCAR3, GPR153, and GPR137 knockdown in wound closure. Our findings indicate that HCAR3 knockdown had the most profound impact, causing a marked reduction in wound closure in primary and N/TERT2G human keratinocytes (Fig. 4A, B, C, and D; S4A, B, and C). siRNA for HCAR3 reduced keratinocyte movement, as measured by accumulated distance travelled by individual cells during the scratch-wound assay (Fig. 4E). Conversely, HCAR3 overexpression resulted in increased keratinocyte movement (Fig. 4F). Supporting the role for HCAR3 in keratinocyte migration, we observed similar effects of HCAR3 knockdown and overexpression in non-confluent cell cultures (Fig. 4G, H, I and J).

**Figure 4:**
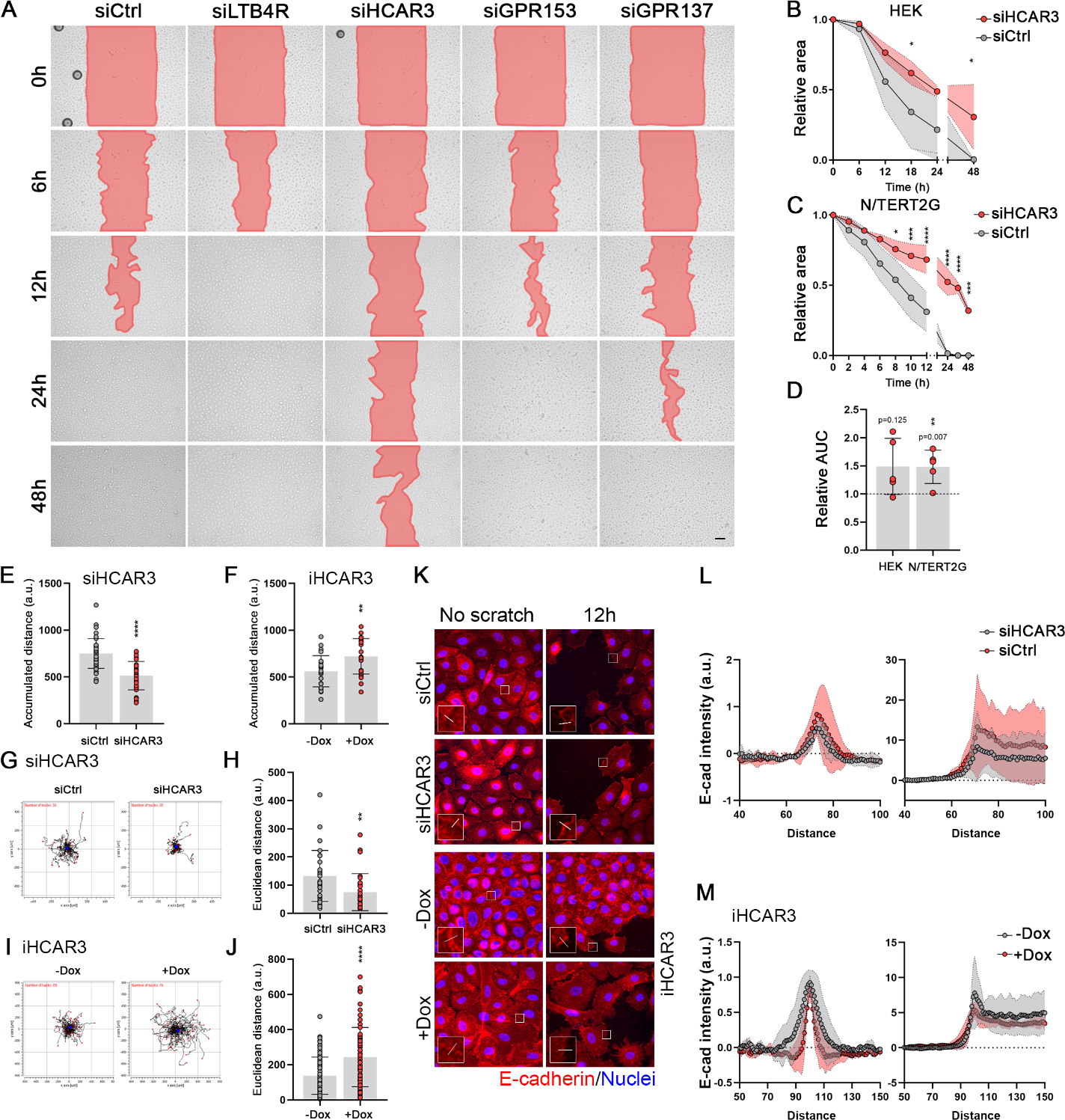
HCAR3 expression affects keratinocytes migration. **(A)** Representative images from scratch assays in N/TERT2G cells transfected with the specified siRNAs for 72hs. The open area is indicated in red color. Scale bar: 100µm. **(B)** and **(C)** Quantification of scratch assays in HEK (B) and N/TERT2G (c) cells. The scratch area was quantified at the indicated times and normalized to time zero. n=5 independent experiments performed in triplicates. **(D)** Quantification of the area under the curve (AUC) for graphs shown on B and C. Data was normalized to siCtrl represented as a black dotted line. n=5 independent experiments performed in triplicates **(E)** and **(F)** Quantification of accumulated distance for individual N/TERT2G cells transfected with the specified siRNAs in the scratch assay (E) or iHCAR3 cells treated as indicated (F). Cells were tracked for 24h. n=24-40 cells per condition. **(G)** and **(H)** Spider plots (G) and quantification of migration expressed as Euclidian distance (H) for individual N/TERT2G cells transfected with the specified siRNAs. Cells were tracked for 24hs. In spider plots, the path of each cell and its final position are represented by a black line and a red dot, respectively. Blue dots are the center of mass. n=30-78 cells per condition. **(I)** and **(J)** Spider plots (I) and Euclidian distance (J) for individual iHCAR3 cells treated as indicated. n=30-78 cells per condition. **(K)** Representative images of E-cadherin expression in confluent cells (no scratch) or 12hs after scratching. E-cadherin is shown in red, and nuclei in blue. Insets show an amplified image of the area marked with a white square. White lines indicate where membrane localization of E-cadherin was quantified. Scale bars: 10 µm. **(L)** and **(M)** Quantification of K. Profile plots show the fluorescence intensity of E-cadherin along a dotted line as indicated in K. Peak of the graph indicates the membrane. Data were normalized to the first value along the quantification lines shown as a black dotted line. N= 25 cells quantified from 3 independent experiments. In B, C, H, and J: no asterisk p>0.05; *p<0.05; **p<0.01; ***p< 0.001; ****p< 0.0001; B, C, and D: 2-way ANOVA followed by t-test; H and J: unpaired t-test.

The scratch-wound assay data suggested that HCAR3 may affect cell-to-cell interaction by influencing the detachment of cells from the monolayer (Movie S1 and 2). Further analysis revealed that while HCAR3 knockdown did not alter total levels of E-cadherin, it affected its membrane localization (Fig. 4K and L, S4D). In contrast, HCAR3 overexpression led to a reduction of membrane E-cadherin (Fig. 4M). Our study highlights a previously unrecognized role for HCAR3 in regulating keratinocyte migration and cell-to-cell attachment.

### HCAR3 knockdown limits keratinocyte respiratory capacity

So far, our results indicate that HCAR3 has a broader impact on keratinocyte biology than the other examined GPCRs. HCAR3 is particularly intriguing because it is expressed only in higher primates, and its functions are not well studied (21). To fill this gap, we focused on understanding the mechanisms by which HCAR3 regulates keratinocyte biology. Since HCAR3 is a Gαi-coupled receptor (22), we investigated whether HCAR3 effects could be recapitulated by Gαi knockdown. Similar to the effects of siHCAR3, siRNA for Gαi reduced keratinocyte proliferation and migration (Fig. S5A, B, C, and D). Gαi has been shown to interact with β-arrestins to regulate migration (23). However, the knockdown of arrestins did not affect keratinocyte proliferation or migration (Fig. S5E, F, G, and H). Our results indicate that HCAR3 might partially regulate keratinocyte biology by activating Gαi but not β-arrestin intracellular signaling pathways.

Given the known role of HCARs in metabolic regulation (22), we hypothesized that HCAR3 might also influence keratinocyte proliferation and migration by affecting cellular metabolism. We analyzed gene networks related to metabolism and mitochondria functions in response to HCAR3 knockdown to test this possibility. We found that siHCAR3 resulted in an upregulation of processes related to fatty acid metabolism and a downregulation of amino-acid metabolism and mitochondrial-related processes (Fig. 5A).

**Figure 5:**
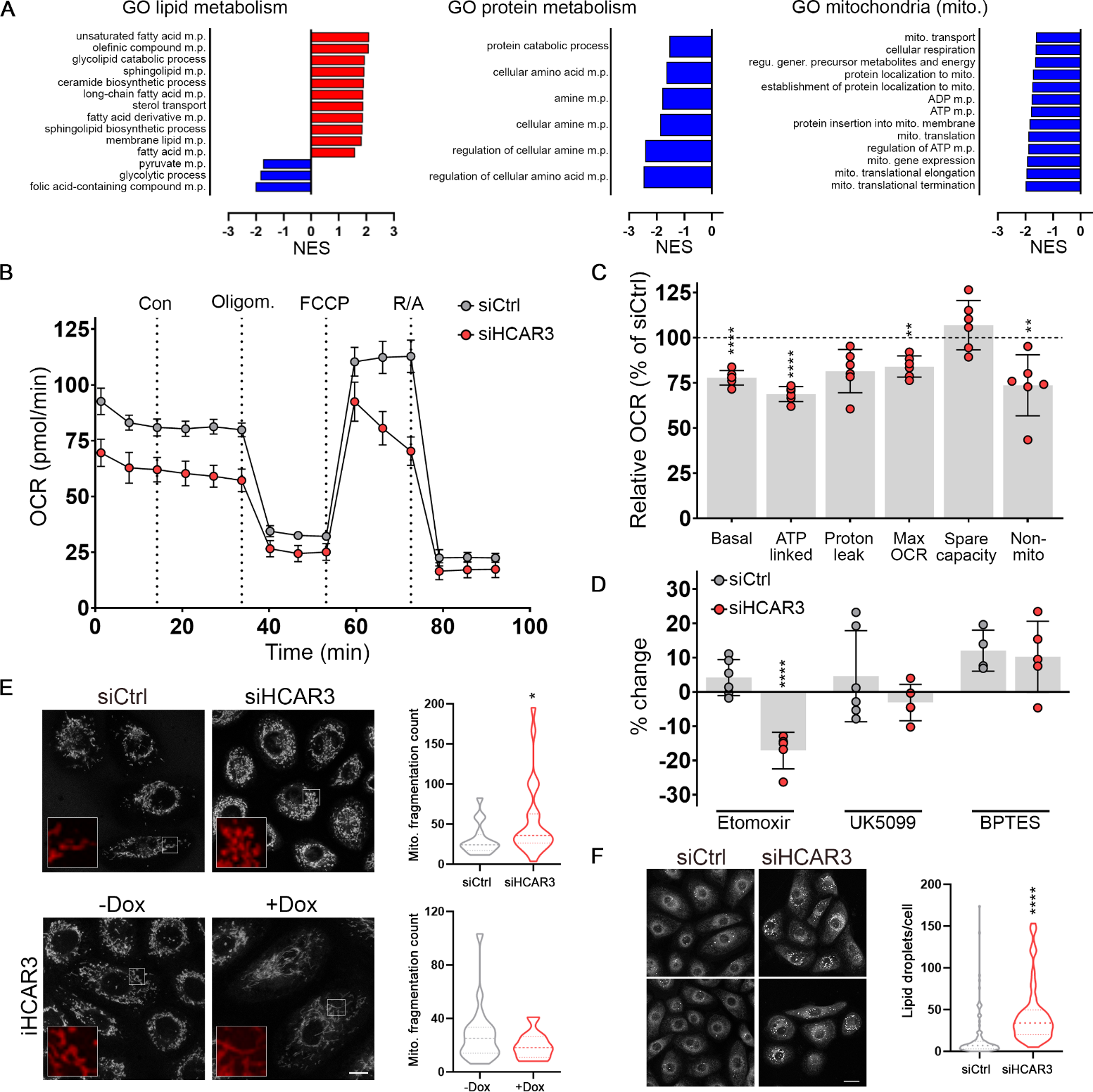
HCAR3 regulates keratinocyte metabolism. **(A)** Graphs of GSEA analysis showing the normalized enrichment score for selected GO biological process in siHCAR3 differentially regulated genes. m.p.: metabolic process, mito.:mitochondria **(B)** Oxygen consumption rates (OCR) normalized to cell number in N/TERT2G cells transfected with a non-targeting siRNA (siCtrl) or siHCAR3 under basal conditions and in response to the indicated drugs (Con: control, Oligom.: oligomycin, R/A: rotenone/antimycin). n= 4-6 replicates. The graph is representative of 2 independent experiments. **(C)** Diagram showing the OCR values calculated from B. **(D)** Percent change in OCR from drug-treated cells with respect to control cells. n= 4-6 replicates. **(E)** Representative images and quantification of mitochondrial staining with MitoTracker in N/TERT2G cells transfected with the specified siRNAs, or iHCAR3 treated as indicated for 72hs. Insets show an amplified image of the area marked with a white square. Violin plots represent the distribution of the mitochondrial fragmentation count per cell. n=9-24 cells per condition. Scale bar: 20 µm. **(F)** Representative images and quantification of lipid droplet staining with LipidTOX in N/TERT2G cells transfected with the specified siRNAs. Violin plots represent the distribution of the number of lipid droplets per cell. n=70-300 cells per condition. Scale bar: 20 µm. In C, D, E, and F: no asterisk p>0.05; *p<0.05; **p<0.01; ****p< 0.0001; C, ANOVA followed by t-test; D, ANOVA followed by Sidak’s multiple comparison test; E, Mann-Whitney test; F, unpaired t-test

To examine the role of HCAR3 in keratinocyte metabolism, we measured oxygen consumption and substrate preference in siHCAR3 cells using a Seahorse Analyzer. We found a drop in the oxygen consumption rate, indicative of a reduction in oxidative phosphorylation (OXPHOS), following HCAR3 knockdown (Fig. 5B and C). We next evaluated the effect of siHCAR3 on metabolic substrate by measuring oxygen consumption in the presence of inhibitors of fatty acid oxidation (etomoxir), pyruvate metabolism (UK5099), or glutamine metabolism (BPTES). Knockdown of HCAR3 increased the reliance on fatty acids for OXPHOS, as evidenced by the decrease in respiratory capacity upon etomoxir treatment in siHCAR3 but not in control cells (Fig. 5D). These results suggest that HCAR3 may regulate the balance of metabolic substrates utilized by keratinocytes, with knockdown of HCAR3 shifting cells towards increased fatty acid utilization. This shift results in energy defects, reducing respiratory capacity and ATP production.

Consistent with the decreased metabolic state of keratinocytes, siHCAR3 caused mitochondrial fragmentation (Fig. 5E), often associated with metabolic stress (24). In contrast, HCAR3 overexpression increased mitochondrial length and connectivity (Fig. 5E). Moreover, siHCAR3 led to an increased number of lipid droplets in keratinocytes (Fig. 5F), further supporting a potential role for this receptor in lipid metabolism. Overall, our findings demonstrate that HCAR3 plays a critical role in regulating gene networks that balance keratinocyte metabolism and mitochondrial morphology to ensure optimal energy outputs.

## Discussion

GPCRs heavily influence epidermal stem cell biology, enabling cells to sense extracellular cues and adapt to environmental changes, maintaining tissue homeostasis (11). To better understand the specific GPCRs that regulate epithelial cells, we used primary cultures of human keratinocytes and identified LTB4R, HCAR3, GPR153, and GPR137 as key receptors that regulate epithelial cell proliferation and differentiation. Furthermore, we determined the transcriptional and signaling networks affected by these GPCRs. Our findings provide valuable insights into the complex interplay between GPCRs and epithelial cell fate decisions.

This study shows that LTB4R, GPR153, and GPR137 affect keratinocyte proliferation and differentiation; however, knockout mice for the homologous genes *Ltb4r1*, *Gpr153,* and *Gpr137* do not have any apparent defects in skin development (14, 25). The lack of an epidermal phenotype in knockout mice shows that these receptors might not be essential for skin homeostasis, although our results suggest that they could have functions during pathological conditions. Indeed, *Ltb4r1* knockout mice have reduced epidermal cell proliferation in response to allergic skin inflammation (14), indicating that LTB4R can drive keratinocyte proliferation in vivo. Similarly, GPR137 could have a role in hyperproliferative diseases since it can regulate cell growth by interacting with Rag-mTORC1 signaling and induce proliferation in cancer cells (26, 27). The overexpression of LTB4R, GPR137, and HCAR3 in 3D organotypic cultures resulted in abnormal skin development and differentiation, suggesting potential pathological effects due to their upregulation.

Proper coordination of metabolic pathways with nutritional requirements is essential to sustain and restore skin homeostasis after injury (28, 29). While this highlights the need for epithelial cells to fine-tune their metabolic state, little is known about how keratinocytes integrate environmental signals into intracellular metabolic pathways. GPCRs can mediate the signaling of numerous metabolites (30), and the HCAR protein family members regulate cellular metabolism under changing nutrient conditions (22). Remarkably, HCAR2 and HCAR3 are some of the most expressed GPCR genes in human keratinocytes (Fig. 1B). HCAR2 (also known as GPR109A) is activated by the antidyslipidemic drug nicotinic acid and has anti-inflammatory and potential anti-psoriatic effects in the skin (31, 32). Although the roles of the closely related receptor HCAR3 are not well known due to its exclusive expression in higher primates (21), our study shows that HCAR3 can affect multiple aspects of epithelial cell biology, including proliferation, differentiation, and migration.

In adipocytes, HCAR receptors are part of a negative feedback loop that senses intermediates from fatty acid β-oxidation in the extracellular space and blocks intracellular lipolysis (22). This information suggests that HCAR3 could have a similar metabolic role in keratinocytes. Indeed, HCAR3 knockdown reduced OXPHOS, possibly by altering metabolite use and mitochondrial morphology. HCAR3 could affect lipid oxidation and de novo lipogenesis in keratinocytes since HCAR3 knockdown caused lipid droplets to accumulate in cells. HCAR2 can function similarly in other cell types (33, 34), indicating that these receptors regulate multiple aspects of lipid metabolism.

The metabolic switch caused by HCAR3 knockdown led to reduced mitochondrial activity and energy output. Interestingly, proliferating keratinocytes are highly dependent on glucose, and reduction of glucose metabolism can lead to defects in proliferation and migration (35). HCAR3 depletion may disturb the metabolic sensing state of keratinocytes, leading to metabolic imbalances between glycolysis and OXPHOS that result in defects in cell proliferation, differentiation, and migration. Our study highlights the importance of proper coordination between extracellular metabolite signaling and intracellular metabolism in maintaining epithelial cell homeostasis and adapting to stress conditions. Further research into the role of metabolite sensing in epithelial cell fate decisions is needed, offering new possibilities for therapeutic interventions for pathological skin conditions and wound healing.

## Materials and methods

### DNA constructs

Codon-optimized sequences for human *LTB4R* (NP_858043.1), *HCAR3* (NP_006009.2), *GPR153* (NP_997253.2) and *GPR137* (NP_001164351.1) with N-terminal 2xHA tag followed by a GDPPVAT linker were purchased from IDT as gBlocks Gene Fragments and cloned in pInducer20 lentiviral inducible vector. pInducer20 was a gift from Stephen Elledge (Addgene plasmid # 44012). Lentiviruses were produced by transfecting Lenti-X 293T cells with pMD2.VSVG (Addgene plasmid # 12259) and psPAX2 (Addgene plasmid # 12260).

### Cell culture and transfections

All cells were cultured at 37°C in the presence of 5% CO2. Lenti-X 293T cells were obtained from Takara Bio and cultured in DMEM (Sigma-Aldrich) containing 10% fetal bovine serum (Sigma-Aldrich). Pooled neonatal primary Human Epidermal Keratinocytes (HEK) were obtained from Life Technologies and were used between passages two to five. N/TERT-2G keratinocyte cell line (36, 37) was provided by Dr. Ellen H. van den Bogaard (Radboud University Medical Center, Nijmegen, The Netherlands) and Dr. James Rheinwald (Brigham and Women’s Hospital, Boston, MA, USA). For GPCR expression analysis, HEK cells were cultured in KGM Keratinocyte Growth Medium BulletKit (Lonza). For all other experiments, HEK and N/TERT-2G keratinocytes were cultured in EpiLife media (Life Technologies) supplemented with Human Keratinocyte Growth Supplement (HKGS S0015, Life Technologies). Where indicated, EpiLife media was supplemented with Defined Growth Supplement (EDGS S0125, Life Technologies), a chemically defined supplement. For differentiation culture conditions, HEK and N/TERT-2G were plated at 100% confluency and EpiLife media supplemented with HKGS and 1.5 mM CaCl_2_. Inducible cell lines were incubated for the indicated time with 0.2 mg/ml of doxycycline to trigger GPCR expression. N/TERT2G cells transduced with inducible expressing GPCRs were used to perform human epidermis reconstruction experiments as previously described (20). The expression of the receptors was confirmed by Western blot or immunofluorescence. For siRNA experiments, cells were transfected with the corresponding siRNAs 24 hours after plating and were analyzed 72 hs after transfection. siRNAs were obtained from Dharmacon/ Horizon Discovery; ON-TARGET plus SMARTpool: ADRB2 (L-005426-01), CELSR2 (L-005460-00), F2RL1 (L-005095-00), ADGRG1 (L-004552-00), LTB4R (L-005653-00), HCAR3 (L-005628-02), HCAR2 (L-006688-02), S1PR1 (L-003655-00), GPRC5A (L-004576-00), LTB4R2 (L-005654-00), S1PR5 (L-004791-00), GPR153 (L-027388-00), F2R (L-005094-00), GPR137 (L-005439-01), LPAR2 (L-004602-00), FZD8 (L-003962-00), ADGRE5 (L-003638-00), FZD7 (L-003671-00), GPR87 (L-005607-00), FZD2 (L-005501-00), FZD6 (L-005505-00), FZD10 (L-005500-00), GPR107 (L-005529-00), TPRA1 (L-005746-00), GNAI1 (L-010404-00), GNAI2 (L-003897-00), GNAI3 (L-005184-00), Non-targeting Control (D-001810-10); siGENOME SMART pool: ARRB1 (M-011971-01), ARRB2 (M-007292-00); siGENOME four individual siRNAs: LTB4R (MQ-005653-01), HCAR3 (MQ-005628-03), GPR153 (MQ-027388-01), GPR137 (MQ-005439-01), ADGRG1 (MQ-004552-00), FZD2 (MQ-005501-01). Cells were transfected with each siRNA at a 5 pmol/well for 96-well plates and 15 pmol/well for 6-well plates, using Lipofectamine RNAiMAX (Invitrogen) according to the manufacturer’s instructions.

### Quantitative PCR

Cells treated as indicated were lysed using the Precellys Lysing Kit (Bertin Instruments). RNA was isolated and processed using RNeasy Plus Mini Kit (Qiagen) according to the manufacturer’s instructions. 1μg of RNA was used to synthesize cDNA with SensiFAST cDNA Synthesis Kit (Bioline) and used as a template for quantitative polymerase chain reaction with reverse transcription (qRT–PCR) analysis using SensiFAST SYBR Lo-ROX Kit (Bioline). Samples were analyzed using a QuantStudio 3 Real-Time PCR System (ThermoFisher). Values were normalized across samples using *RPLP0* expression. Oligonucleotides used for amplification were (Gene, Forward sequence, Reverse sequence): RPLP0, 5’-TGTCTGCTCCCACAATGAAAC-3’, 5’-TCGTCTTTAAACCCTGCGTG-3’; ADRB2, 5’-CGCTACTTTGCCATTACTTCAC-3’, 5’-GACACAATCCACACCATCAGA-3’; CELSR2, 5’-CCTGCCTCTGTCGTGATG-3’, 5’-GCAATCGCACTTGAAACCG-3’; F2RL1, 5’-CGTTCTTTGCATGATCCCTGA-3’, 5’-GCCATGTCTATGCCCTGTA-3’; ADGRG1, 5’-GTTAGCCACTTCTGTCACCTC-3’, 5’-CTTTTGAGCTCGCACATGTC-3’; LTB4R, 5’-TGCAGAAGATGTAGTGTTCATGG-3’, 5’-CCTTTGACAGCAGACCCTAC-3’; HCAR3, 5’-<colcnt=5> GACATTACTCGATGCAACAGC-3’, 5’-GGCCCAACCTCAAATAACCA-3’; HCAR2, 5’-CCCTGAGTCCATTTCTGCTAA-3’, ACGATAATTACGGTCTTGAGCA-3’, 5’-GATTGGAGGGAATTGTGTTGC-3’, 5’-GCAGAATCTACTCCTTGGTCA-3’; S1PR1, GPRC5A, 5’-5’-GAAGCAGCACCAAGTTCAC-3’, 5’-GTACTTGGATTTCAGGCCATTG-3’; LTB4R2, 5’-GTAGCAGACCGACATCCTTC-3’, 5’-GGACAGGAGTAGGCACCT-3’; S1PR5, 5’-TAATGCAGGACGATGACCTC-3’, 5’-AAGCAACCCACTGAAGACC-3’; GPR153, 5’-CACCACAGCACGCAGAG-3’, 5’-CTCGTGACCACCATAGTCTTC-3’; F2R, 5’-GGCCAGACAAGTGAAGGAA-3’, 5’-CGCCTCTATCTTGCTCATGAC-3’; GPR137, 5’-CAGCAGAAAGAGCAGCGA-3’, 5’-CACCTTGACGCTTATGAACCT-3’; LPAR2, 5’-<colcnt=4>GTACTTTTCTACAGCCAGGACA-3’, 5’-AGCCTGGTCAAGACTGTTG-3’; FZD8, 5’-GAATACCCAAACCGTGCAATC-3’,5’-TTTCTCCCTCGACTCTTCCTA-3’;ADGRE5,5’-CTGTCGCTGTCACTCTTCC-3’, 5’-CCATCCTTTTCGCCTTCTCC-3’; FZD7, 5’-GCAGTTCTTTTCCCTACCGT-3’, 5’-CTACCACAGACTTAGCCACAG-3’; GPR87, 5’-TGGGTTCAGCATAGGTTATTCC-3’, 5’-GCCGACCTTAGTTTCAAAGC-3’; FZD2, 5’-CCCACCCGTCTTTATCACTT-3’, 5’-TTTCCCTGCAGTCTATTCACTC-3’; FZD6, 5’-TGTGCATGTCTTTTTGCTTCC-3’, 5’-TCATTGTCGTCAGTACCATATCC-3’; FZD10, 5’-TCTTCCAGTAATCCATGTTGAGG-3’, 5’-AGAAGCTCATGGTGCGTATC-3’; GPR107, 5’-GAGACTCTATGCGACTTCCTG-3’, 5’-TGCCATCCTCTTCCCAGT-3’; TPRA1, 5’-TCCACTTGGCATTTGTAGGAG-3’, 5’-CCACAACCTTCCTGTACTTCA-3’; GNAI1, 5’-TTAGGGCTATGGGGAGGTTGA-3’,5’-GGTACTCTCGGGATCTGTTGAAA-3’; GNAI2,5’-GACGCCAGGCAGCTATTTG-3’,5’-ACTCTGTGCAATACGCTCCAG-3’; GNAI3,5’-GACGGCTAAAGATTGACTTTGGG-3’,5’-CCGTTTAATCACTCCTGCTAGTT-3’; ARRB1,5’-CAGGAACGCCTCATCAAGA-3’,5’-GCAGTGTCACAGAACATGGA-3’; ARRB2, 5’-AGACAGTACGCCGACATCT-3’, 5’-GATTCCCAGCACCTCCTTG-3’.

### Gene expression analysis

For GPCR expression analysis, primary human keratinocytes were cultured under basal or non-differentiating (low confluency, low calcium) and differentiation (high confluency, high calcium) conditions and harvested after 24h (basal) and 24h, 48h, and 96h (differentiation) in triplicates. For knockdown experiments, cells were plated in 6 well plates and transfected with the corresponding siRNA in triplicate as detailed before; the following day media was changed, and cells were harvested after 72h. RNA was isolated, as desctribed above and RNA integrity was measured with the TapeStation system (Agilent). Samples with RIN>8 were further processed. mRNA expression profiling was performed in the CCR-Sequencing Facility at the NIH. For GPCR expression, reads of the samples were trimmed for adapters and low-quality bases using Trimmomatic software before alignment with the reference genome Human-hg19 and the annotated transcripts using STAR. Gene counts were filtered by genes with ≥5 reads and normalized to TMM (Trimmed Mean of M values) and TPM (Transcripts Per Million) using PartekFlow software, version 7.0 (Partek Inc). TMM normalized counts were used for differential analysis using the PartekFlow GSA algorithm (Partek Inc). GPCR information and coupling was obtained from GPCRdb (38) and GPCRs were selected based on TPM≥0.5 and changes during differentiation were selected based on q<0.05 and fold change ≥1.5 for upregulated and ≤-1.5 for downregulated. A circular plot for GPCR expression was created using ClicO FS (39). For siRNA experiments, counts were calculated in PartekFlow v8 and processed with R version 4.0.3 using the edgeR 3.31.4 R package. Batch effects from different sequencing experiments were removed using the removeBatchEffect R function from the limma 3.45.14 R package using siCon samples as reference. Differential expression analysis against siCon cells was performed using the limma 3.45.14 R package. Gene ontology terms and normalized enrichment scores (NES) in individual datasets were calculated using fgsea 1.16.0 R package. Differentially expressed genes were selected based on adjusted p value<0.05 and fold change ≥1.5 for upregulated and ≤-1.5 for downregulated genes. Gene ontology terms in overlapping genes were calculated using ToppGene (40). Upstream regulators analysis was generated using Ingenuity Pathway Analysis (IPA, Ingenuity Systems, www.ingenuity.com). Differentiation and signaling pathways gene lists were modified from (41).

### Immunoblot analysis

Western blot assays were performed as described previously (20). Antibodies used were: K10 (BioLegend; catalog no. 905404; 1:200), KLF4 (Cell Signaling; catalog no. 12173S; 1:1000), pERK (Cell Signaling; catalog no. 4370; 1:1000), ERK (Cell Signaling; catalog no. 4695; 1:1000), pAKT(Thr308) (Cell Signaling; catalog no. 13038; 1:1000), pAKT(Ser473) (Cell Signaling; catalog no. 4060; 1:1000), AKT (Cell Signaling; catalog no. 4691; 1:1000), GAPDH (Cell Signaling; catalog no. 2118; 1:2000), pS6 (Cell Signaling; catalog no. 2211; 1:1000), S6 (Cell Signaling; catalog no. 2211; 1:1000), CREB (Cell Signaling; catalog no. 9197; 1:1000), pCREB(Ser133) (Cell Signaling; catalog no. 9198; 1:1000), E-cadherin (Cell Signaling; catalog no. 3195S; 1:1000). The secondary antibody used was Anti-rabbit IgG HRP linked (Cell Signaling; catalog no. 2217; 1:5000). Bands were detected using a ChemiDoc™ Imaging System (Bio-Rad) with Clarity™ Western ECL Blotting Substrates (Bio-Rad) according to the manufacturer’s instructions.

### Immunofluorescence and quantification

For immunofluorescence, cells were fixed with 3.2% paraformaldehyde in PBS, permeabilized with 0.05% Triton X-100 and 200 mM glycine in PBS and blocked with BSA 3% in PBS. Cells were incubated with the primary antibodies overnight at 4°C, followed by two hs incubation with the secondary antibodies. The following antibodies were used: E-cadherin (Cell Signaling; catalog no. 3195S; 1:500), HA-Tag (Cell Signaling; catalog no. 3724; 1:500); K10 (BioLegend; catalog no. 905404; 1:200); Donkey anti-Rabbit IgG Alexa Fluor 555 (Invitrogen; catalog no. A-31572; 1:1000). Nuclei were stained with Hoechst 33342 (Invitrogen). Immunofluorescence and histology analysis of the organotypic tissues was performed on tissue sections embedded in paraffin and stained as previously described (20). Images were obtained using either a Leica SP8 confocal microscope with LASX software or a Keyence BZ-X700 with automatic stage and focus with BZX software. The final images were bright contrast adjusted with Adobe Photoshop. Quantification of plasma membrane E-cadherin was done in ImageJ using the plot profile tool.

### Cell proliferation

Cells were plated in 96 well plates and transfected the next day, as explained above. For differentiated conditions, and Ca^2+^ was added to the media the following day after plating. 72hs after transfection, cell proliferation was evaluated using 4hs incorporation of EdU followed by labeling with the Click-IT EdU Imaging Kit (Invitrogen) according to the manufacturer’s instructions. Images were obtained using a Keyence BZ-X700 with automatic stage and focus with BZX software. The final images were bright contrast adjusted with Adobe Photoshop. Quantification of proliferative cells was done in the BZX analysis software (Keyence) equipped with hybrid cell count and macro cell count. Total cells were counted in the Hoechst 33342 channel (nuclei), and EdU-positive cells were counted in the corresponding channel. Values shown are the proportion of positive cells/total cells per field normalized to a control condition. Between 500 to 1000 cells per field were quantified in 9 different areas per condition in 3-4 independent experiments performed by triplicate. Proliferation score for pooled versus individual siRNAs was calculated as described in (26) using the formula (x-µ)/sd, where x is the average of the raw values for each experiment, µ is the mean of the proliferation value using a non-targeting siRNA as control (µ=1), and sd is the standard deviation of each experiment performed in triplicates. The scores were the average for the four independent experiments. Positive values correspond to an increase in proliferation, and negative results correspond to a reduction in the proliferation rate. Individual siRNAs having a similar behavior to pooled siRNAs reduce the possibility of off-target effects.

### Scratch assay

To study cell migration, HEK or N/TERT2G cells were plated in 96-well plates and transfected 24 hs later with the indicated siRNAs. 72 hrs after transfection, with 100% confluent cells, a scratch was made using a wound maker system (Essen BioScience) following the manufacturer’s indications. Data were collected every 15 minutes up to 6 hours to 24 hours using a Keyence BZ-X700 microscope equipped with a CO2 and temperature control chamber. The wound area was quantified, and videos created using ImageJ.

### Mitochondria bioenergetic analysis

A Seahorse XFe96 Analyzer and XF Mito Stress Test Kit (Agilent Technologies) were used to measure the oxygen consumption rate (OCR) in N/TERT2G cells with differential expression of HCAR3. Cells were seeded and transfected with siHCAR3, as explained above. 24h before the experiment, cells were plated onto an XF96 Cell Culture Microplate (Agilent Technologies) at 4×10^4^ cells/well. The day before the assay, the cartridge sensor was hydrated overnight with Seahorse Bioscience XF24 Calibration Buffer at 37 °C without CO2. On the day of the assay, media was replaced with serum-free Seahorse XF base media, and cells were incubated at 37 °C in a non-CO2 incubator for 1h. OCR was measured after the injection of oligomycin (1.5 µM), FCCP (2 µM), and rotenone/antimycin (0.5 µM). In addition to XF Mito Stress Test Kit, XF Mito Fuel kit (Agilent Technologies) was used together that includes three inhibitors, UK5099 (2 µM), BPTES (3 µM) and etomoxir (4 µM) to measure the dependency of cells to the three major mitochondrial fuels: glucose, glutamine and long-chain fatty acids, respectively. Inhibitors were added at the control injections time shown in Fig. 5B. Data were normalized to the number of cells per well and analyzed using Agilent Seahorse Analytics. For each assay, 4-6 technical replicates were measured. Equal plating was confirmed after the assay by measuring the total protein per well.

### Lipid droplet staining

HCS LipidTOX Green Neutral Lipid Stain (Thermo Fisher Scientific) was used to analyze lipid droplets following the manufacturer’s indications. Briefly, cells were plated and transfected as described above. On the day of the experiment, cells were fixed with 3.2% paraformaldehyde and incubated for 30 minutes at room temperature with LipidTOX (1:1000) in PBS. Cells were analyzed using a Keyence BZ-X700 microscope without washing. Images were quantified using ImageJ.

### Mitochondria staining

Cells were plated in a Nunc Lab-Tek Chambered Coverglass (Thermo Fisher Scientific) and transfected or treated with doxycycline as described above. On the day of the experiment, cells were incubated for 30 minutes with 100 nM MitoTracker (Thermo Fisher Scientific) at 37°C. Cells were washed and prepared for analysis by microscopy. Mitochondrial fragmentation was quantified using ImageJ as described in (42).

### Statistical analysis

Statistical analyses, variation estimation, and validation of test assumptions were carried out using the Prism 5 statistical analysis program (GraphPad). Asterisks denote statistical significance (non-significant or NS, p > 0.05; *p < 0.05; **p < 0.01; ***p < 0.001; ****p < 0.0001). All data are reported as mean ± standard deviation (SD).

## Data availability

Raw and processed RNAseq data generated for this study are available in the GEO database under accession codes: GSE228631 and GSE228651. *Gpr137* and *Gpr153* knockout phenotype data was retrieved from the www.mousephenotype.org (25). Single cell expression levels for GPCRs in human skin were obtained from v22.proteinatlas.org (19), data available at: https://www.proteinatlas.org/ENSG00000213903-LTB4R/single+cell+type/skin, https://www.proteinatlas.org/ENSG00000255398-HCAR3/single+cell+type/skin, https://www.proteinatlas.org/ENSG00000173264-GPR137/single+cell+type/skin, https://www.proteinatlas.org/ENSG00000158292-GPR153/single+cell+type/skin.

## Author Contributions

MPP, KL, SWSK, CHS and RIB performed experiments; MPP, SWSK, TC and RIB analyzed data and prepared figures; TC and RIB performed bioinformatic analysis; NPS assisted with metabolism studies; MPP and RIB compiled final figures; MPP and RIB wrote the manuscript draft; MPP, KL, CHS, NPS and RIB edited the manuscript.

## Acknowledgments

This research was supported by the Intramural Research Program of the National Institutes of Health, National Cancer Institute, Center for Cancer Research (ZIA BC 011764 and ZIA BC 011763). This work used the computational resources of the NIH High-Performance Computing Biowulf Cluster. We thank the members of the CCR Sequencing Facility at Frederick National Laboratory for Cancer Research for their help during sample preparation, sequencing, and data processing.

## Supplementary Material Legends

**Movie S1:** Representative movie of scratch assay for cells treated with siHCAR3 or non-targeting siRNA (siCtrl). Each frame was acquired every 15 min, movie is 24 hs. Scale bar: 100 µm

**Movie S2:** Representative movie of scratch assay for inducible HCAR3 cells treated or not with doxycycline (DOX) to induce expression of the receptor. Each frame was acquired every 15 min, movie is 24 hs. Scale bar: 100 µm

---

**Figure S1:**
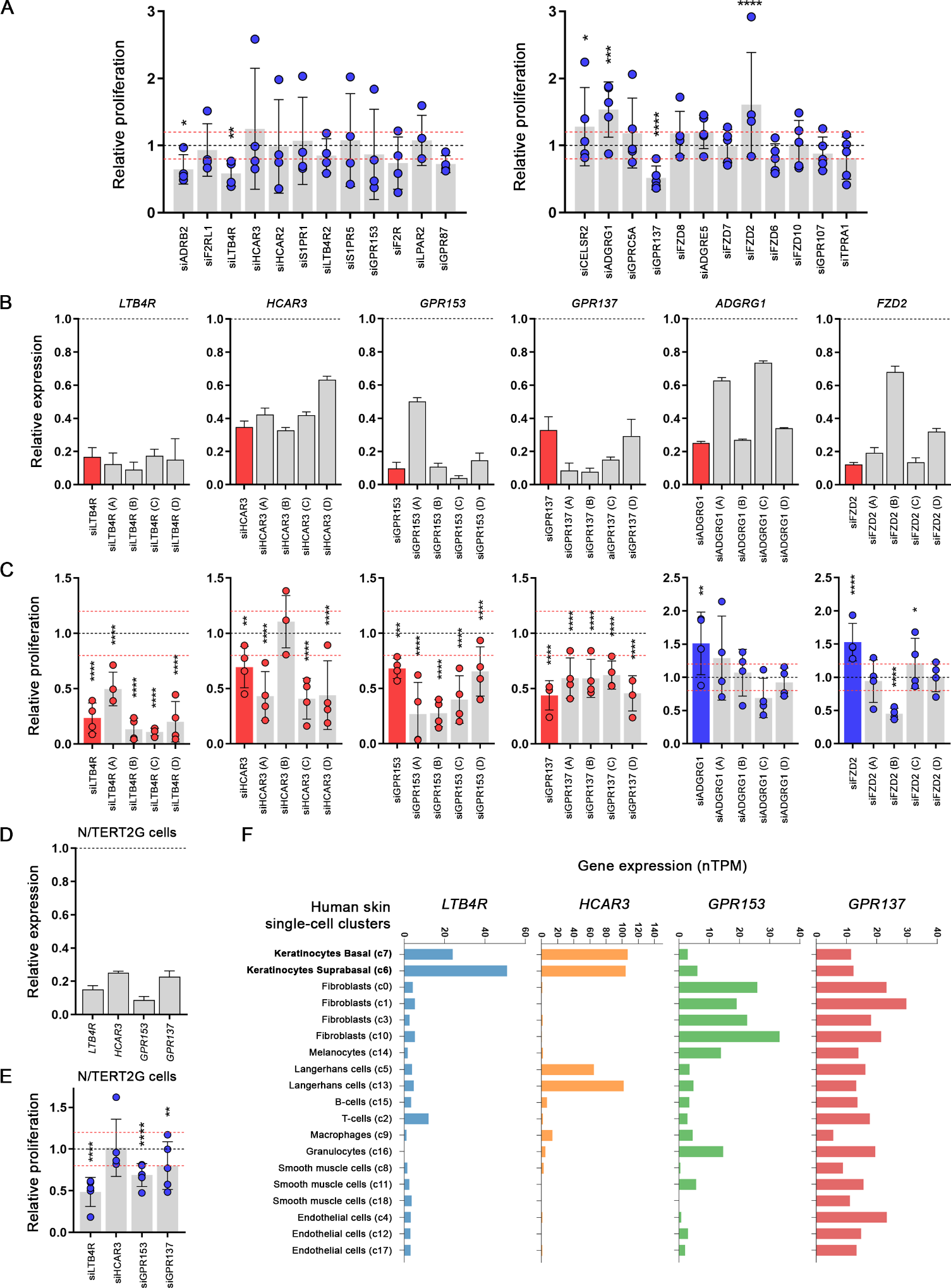
**(A)** Quantification of EdU-proliferation assay performed 72h after siRNA transfection with the indicated GPCRs siRNAs (Class A left panel, non-Class A right panel) in differentiated primary HEK cells (48h differentiation). Results are normalized and compared to a non-targeting siRNA, represented as a black dotted line. The values of ±20% change in proliferation are shown as red dotted lines. n=4-5 independent experiments. **(B)** qRT-PCR analysis showing the expression of the selected GPCRs in HEK cells 72 h after siRNA transfection. Values are indicated as fold over non-targeting siRNA control, shown as a black dotted line. Solid color bar represents pooled siRNA, and gray bars are the results of each individual siRNAs (A-D). **(C)** Quantification of EdU-proliferation assays performed 72h after siRNA transfection with the indicated GPCRs siRNAs in primary HEK cells. Solid color bar represents pooled siRNA, and gray bars are the results of each individual siRNAs. Red indicates the assay was performed in non-differentiated conditions; blue indicates the assay was under differentiated conditions. Results are presented as in A. n=4 independent experiments. **(D)** qRT-PCR analysis showing the expression of the indicated GPCRs in N/TERT2G cells 72h after transfection with the corresponding siRNA. Values are presented as in B. **(E)** Quantification of EdU-proliferation assays in differentiated N/TERT2G cells transfected with the inidicated siRNAs. Results are presented as in A. n=5 independent experiments. **(F)** Expression of the indicated GPCRs as normalized transcripts per million (nTPM) in data from single-cell mRNA expression from human skin. Different cell populations are shown based on their corresponding cluster (indicated as "c") derived from clustering analysis. Data obtained from v22.proteinatlas.org. In A, C and E: no asterisk p>0.05; *p<0.05; **p<0.01; ***p< 0.001; ****p< 0.0001; 2-way ANOVA followed by t-test.

**Figure S2:**
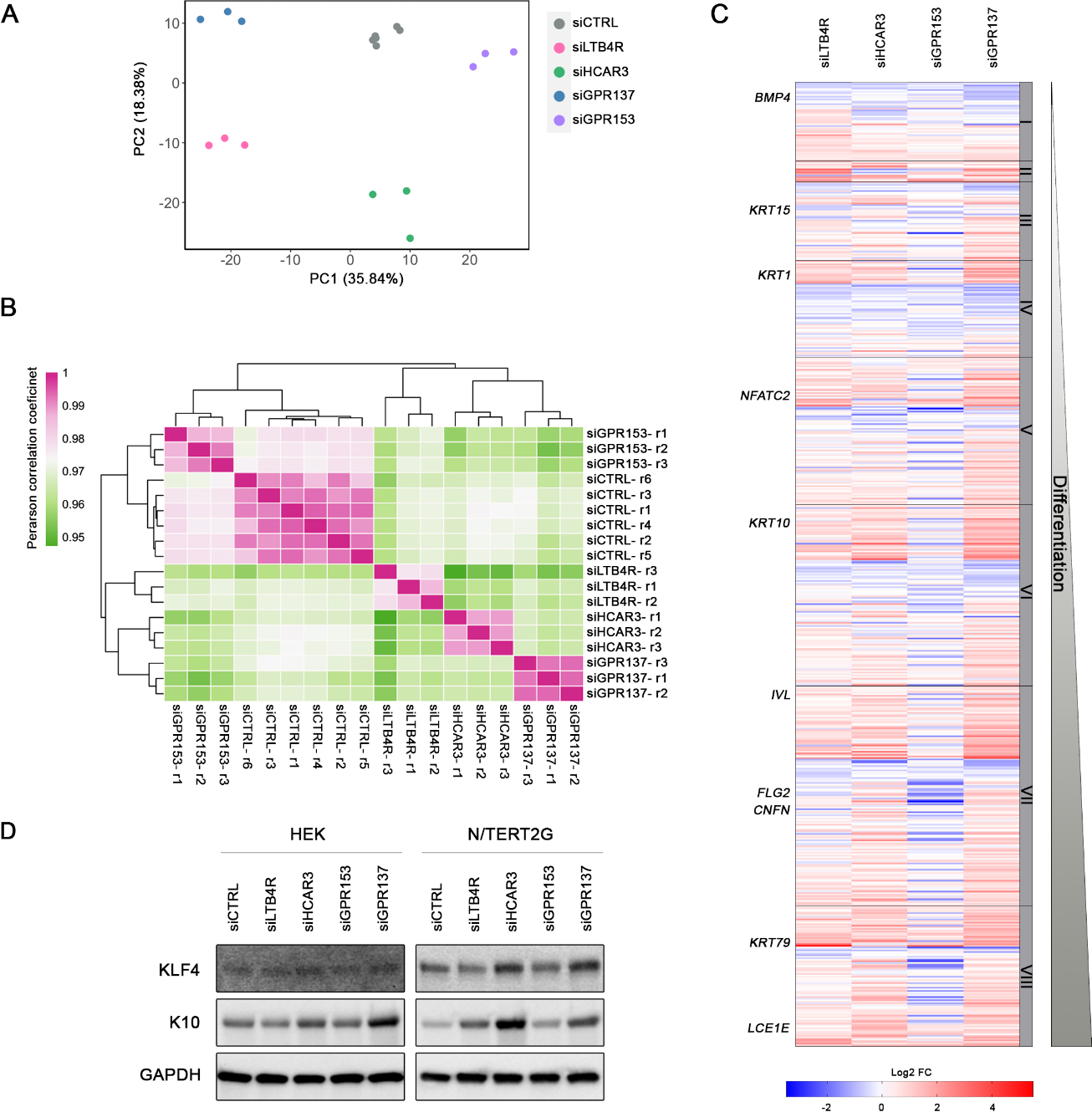
**(A)** Principal component analysis (PCA) showing the results for the RNAseq data in HEK cells transfected with the indicated siRNAs. (n=6 independent samples for siCTRL and n=3 independent samples for each siGPCR). **(B)** Pearson correlation coefficient and clustering among individual samples from the RNAseq experiment. r indicates repetition (independent sample). **(C)** Graph showing the fold change (Log2 FC) of genes expressed in the different layers in the epidermis for each of the siGPCRs datasets. Groups I-VIII contain genes clustered according to their expression pattern during differentiation, with I representing the basal compartment and VIII representing the most differentiated layer. Examples of representative genes are shown to the left. **(D)** Western blot showing keratin 10 (K10) and KLF4 expression in primary HEK or N/TERT2G cells 72h after transfection with the indicated siRNAs.

**Figure S3:**
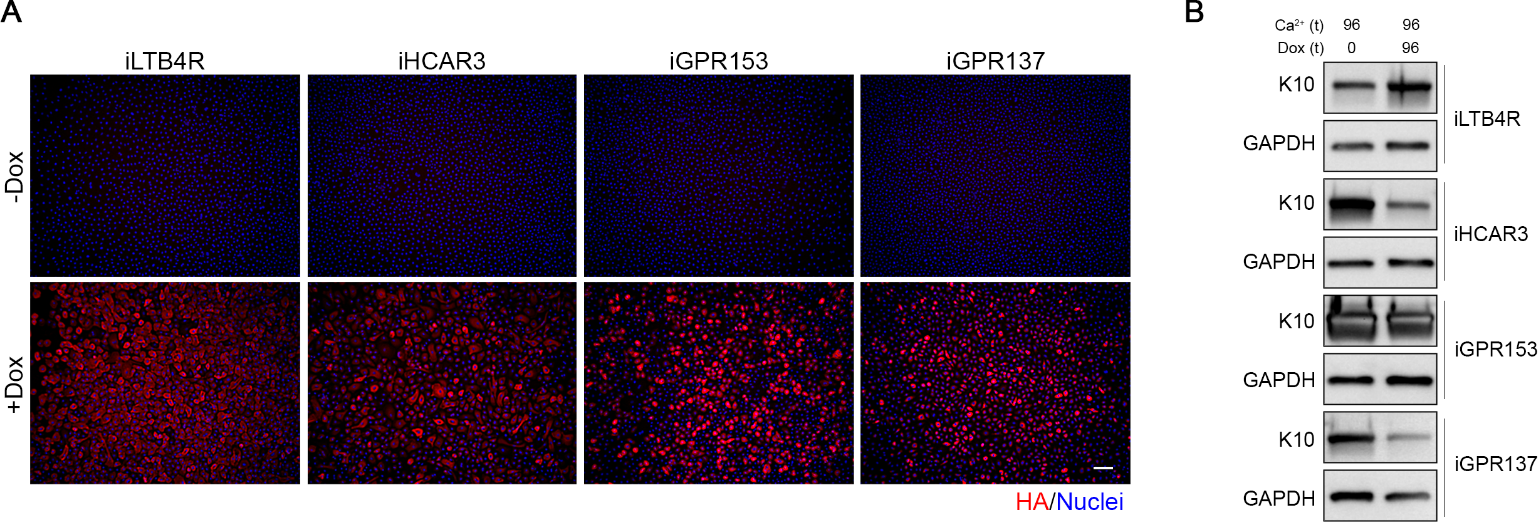
**(A)** Images showing expression of inducible (i) GPCRs in N/TERT2G treated (+Dox) or not (-Dox) with doxycycline for 24 h. GPCRs were tagged with N-terminal HA. HA signal is shown in red, and nuclei are shown in blue. Scale bar: 100µm. **(b)** Western blot showing the levels of Keratin 10 (K10) in N/TERT2G in cells with GPCRs-inducible expression treated (+Dox) or not (-Dox) with doxycycline for 96 h under differentiation conditions.

**Figure S4:**
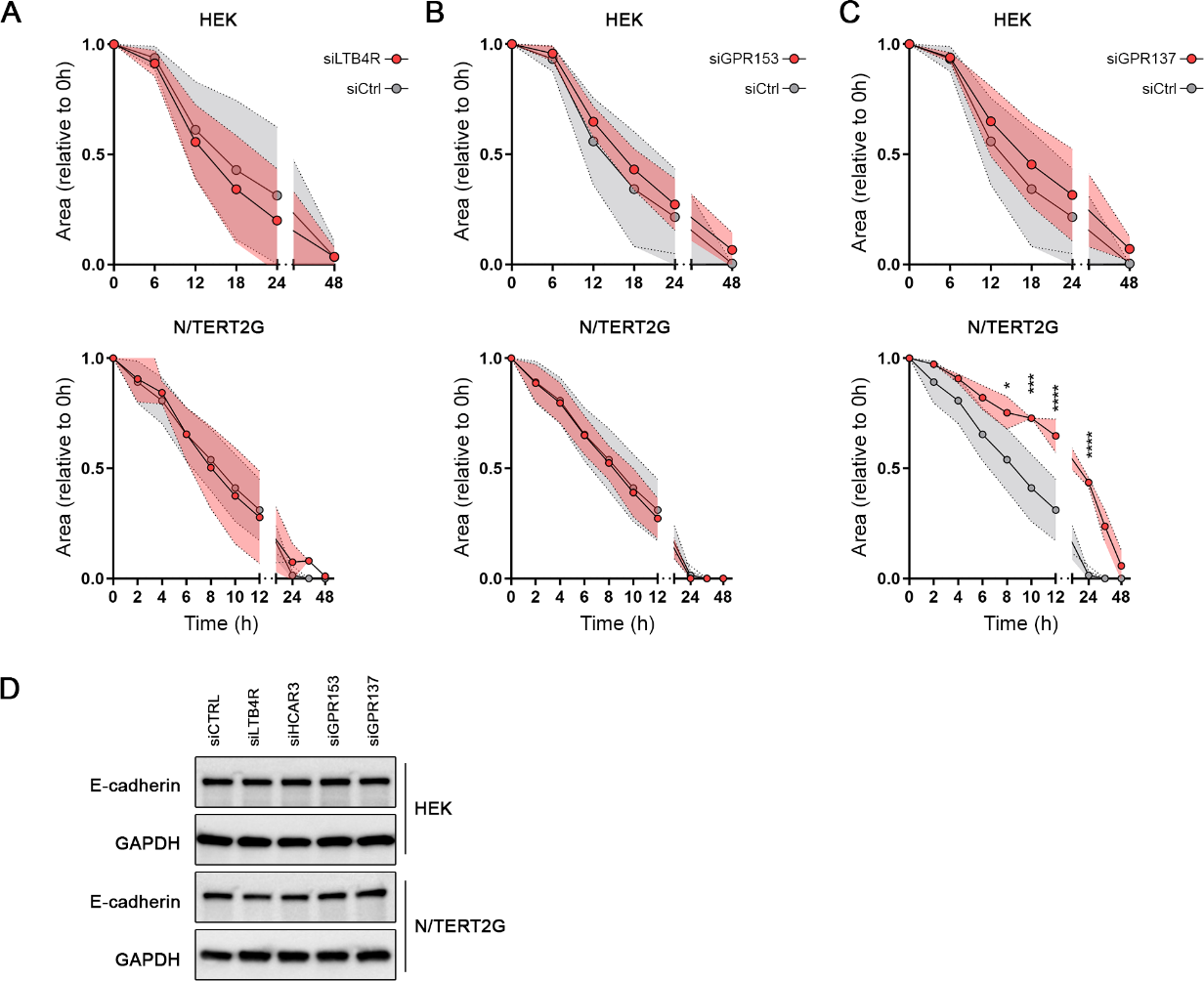
**(A)**, **(B),** and **(C)** Quantification of scratch assays in primary HEK (top) and N/TERT2G cells (bottom) for the indicated GPCRs knockdowns. The scratch area was quantified at the indicated times and normalized to t=0. n= 4-5 independent experiments performed in triplicates. **(D)** Western blot showing the levels of E-cadherin in HEK and N/TERT2G cells 72h after transfection with siRNAs.

**Figure S5:**
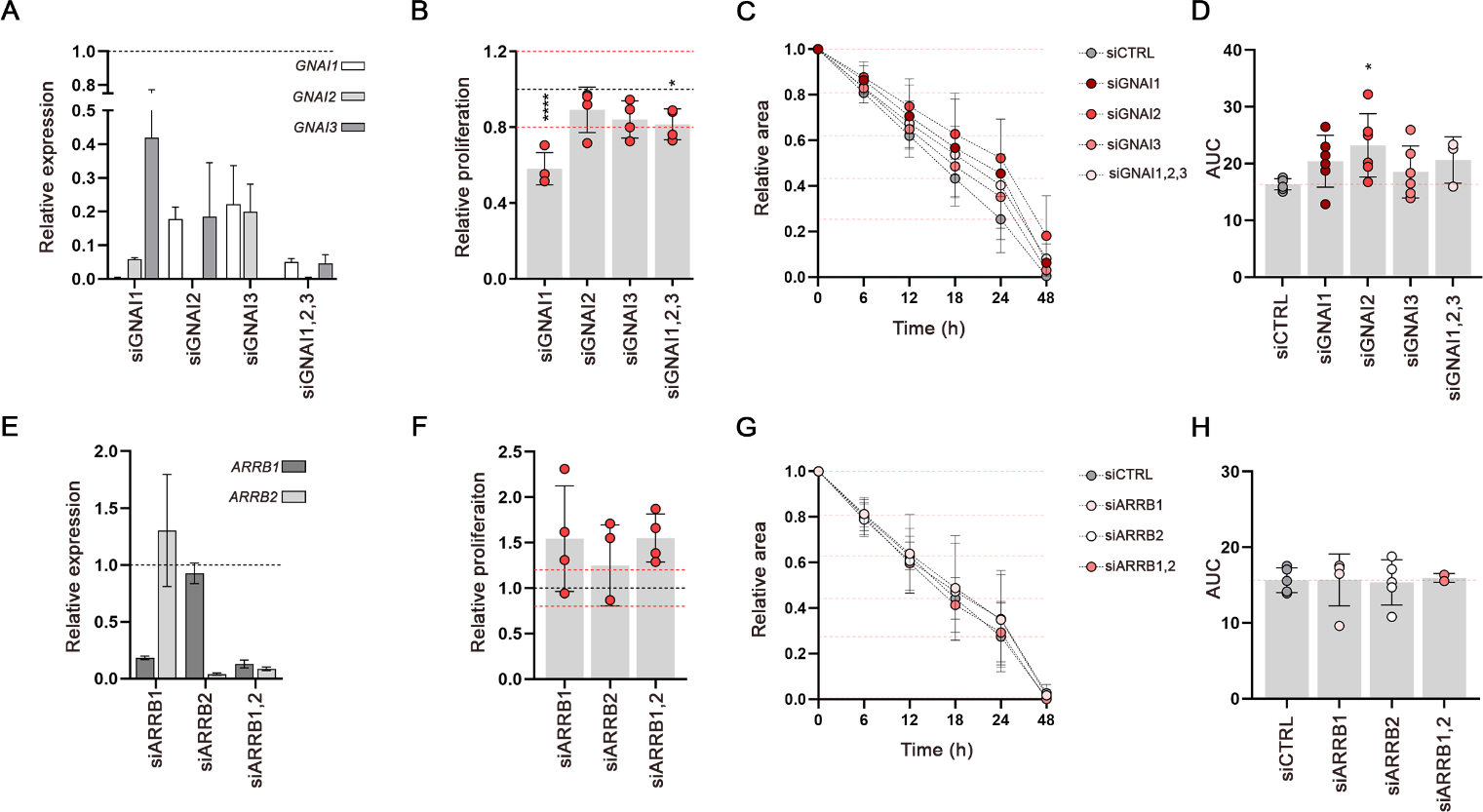
**(A)** qRT-PCR analysis of the indicated genes in N/TERT2G cells 72h after siRNA transfection. Values are displayed as fold over non-targeting siRNA, shown as a black dotted line. **(B)** Quantification of EdU-proliferation assay performed 7 h after siRNA transfection with the indicated siRNAs in N/TERT2G cells. Results are normalized and compared to a non-targeting siRNA, represented as a black dotted line. ±20% change in proliferation is shown as red dotted lines. n=4 independent experiments performed in triplicates. **(C)** Quantification of scratch assays in N/TERT2G cells transfected with the indicated siRNAs for 72h. The scratch area was quantified at the indicated times and normalized to t=0. n= 3-6 independent experiments performed in triplicates. Red dotted lines indicate the values of siCTRL (non-targeting siRNA) at the different time points. **(D)** Quantification of the area under the curve (AUC) from E. **(E)** qRT-PCR analysis showing the expression of indicated genes in N/TERT2G cells 72h after siRNA transfection. Values are displayed as fold-over siCTRL (non-targeting control) transfected cells, shown as a black dotted line. **(F)** Quantification of EdU-proliferation assay performed 72h after siRNA transfection with the indicated siRNAs in N/TERT2G cells presented as in B. n= 4 independent experiments performed in triplicates. **(G)** Quantification of scratch assays in N/TERT2G cells transfected with the indicated siRNAs for 72h presented as in C. n= 2-5 independent experiments performed in triplicates. **(H)** Quantification of the area under the curve (AUC) shown on (G). B, D, F, and H: no asterisk p>0.05; *p<0.05; ****p< 0.0001; B and F: one-way ANOVA followed by t-test; D and H: one-way ANOVA followed by t-test.

